# Targeted demethylation of H3K9me3 and H3K36me3 improves somatic cell reprogramming into cloned preimplantation but not postimplantation bovine concepti

**DOI:** 10.1101/699181

**Authors:** Fanli Meng, Kathrin Stamms, Romina Bennewitz, Andria Green, Fleur Oback, Pavla Turner, Jingwei Wei, Björn Oback

## Abstract

Correct reprogramming of epigenetic marks in the donor nuclei is a prerequisite for successful cloning by somatic cell transfer. In several mammalian species, repressive histone (H) lysine (K) trimethylation (me3) marks, in particular H3K9me3, form a major barrier to somatic cell reprogramming into pluripotency and totipotency. We engineered bovine embryonic fibroblasts for the doxycycline-inducible expression of Kdm4b, a demethylase that removes histone 3 lysine 9 trimethylation (H3K9me3) and H3K36me3 marks. Upon inducing *Kdm4b*, H3K9me3 and H3K36me3 levels reduced ∼3-fold and ∼5-fold, respectively, compared to non-induced controls. Donor cell quiescence has been previously associated with reduced somatic trimethylation levels and increased cloning efficiency in cattle. Simultaneously inducing *Kdm4b* expression (via doxycycline) and quiescence (via serum starvation), further reduced global H3K9me3 and H3K36me3 levels by a total of 18-fold and 35-fold, respectively, compared to non-induced, non-starved control fibroblasts. Following somatic cell transfer, *Kdm4b*-BEF fibroblasts reprogrammed significantly better into cloned blastocysts than non-induced donor cells. However, detrimethylated donors and sustained *Kdm4b*-induction during embryo culture did not increase rates of post-blastocyst development from implantation to survival into adulthood. In summary, KDM4B only improved somatic cell reprogramming into early preimplantation stages, highlighting the need for alternative experimental approaches to reliably improve somatic cloning efficiency in cattle.

## INTRODUCTION

Despite their morphological and functional diversity, different somatic cell types within an individual contain the same genetic information. Phenotypic differences are stabilised by epigenetic modifications, such as histone methylation, that regulate cell-specific gene activity.

‘Reprogramming’ of these epigenetic marks naturally occurs during mammalian gametogenesis and preimplantation embryogenesis [1]. Cloning by somatic cell transfer (SCT), can artificially reprogram somatic cells [2]. During SCT, a somatic donor cell is fused with an enucleated recipient oocyte (or cytoplast). Epigenetic marks on the donor chromatin are then reprogrammed by cytoplast factors that are still largely unknown. In some cases, the donor nucleus regains totipotency, i.e. the ability to form all embryonic and extra-embryonic lineages in a viable animal. However, totipotency reprogramming of somatic cells is inefficient due to a strong resistance to erase the epigenetic memory of previous lineage decisions and restart embryonic gene transcription [3]. This reprogramming resistance compromises development of the SCT embryo, resulting in aberrant methylation patterns of DNA [4] and histones [5] and dysregulation of gene expression [6].

To improve epigenetic reprograming after SCT, several approaches have been used with some success. First, in chronological order of discovery, pharmacological histone deacetylation inhibitors (HDACi) induce hyperacetylated, transcriptionally permissive chromatin and can increase *in vivo* cloning efficiency in mouse [7, 8] and pig [9] but their impact in cattle species remains controversial [10-12]. Second, cloned mouse embryos often overexpress *Xist*, a noncoding gene responsible for X chromosome inactivation, from the active X chromosome, ectopically downregulating many X-linked genes necessary for embryonic development [13]. Normalising *Xist* expression by gene knockout or knockdown can increase cloning efficiency by up to 20% per embryo transferred [13, 14]. Third, we discovered that heterochromatic H3K9me3 marks pose a major epigenetic barrier in mouse cloning, antagonising an open, transcriptionally permissive chromatin configuration [15]. Overexpressing the histone lysine demethylase *Kdm4b* in donor cells decreased H3K9me3 levels in embryonic stem cells [15] and this demethylation was even more pronounced in mouse embryonic fibroblasts (MEFs) [16]. Reduced H3K9me3 donor levels correlated with improved development of embryos cloned from ESCs [15] and, even more markedly, from MEFs [16]. The principle we first described in mouse was confirmed by other groups, using various *Kdm4* isoforms, and extended to other mammalian species. In mouse, *Kdm4b* expression was aberrant in developmentally arrested SCT two-cells, a defect which could be overcome by injecting exogenous *Kdm4b* mRNA into enucleated MII oocytes prior to SCT [17]. Injecting SCT embryos with *Kdm4d* mRNA also markedly increased clone development in mouse [18]. The same approach was then used for overexpressing another related H3K9me3 demethylase, *KDM4A*, to improve blastocyst formation rate in human SCT embryos [19]. In non-human primates, successful cloning of cynomolgus monkeys was also attributed to *KDM4D* mRNA injection [20]. Combined with HDACi treatment, this approach greatly improved blastocyst development and pregnancy rate of transplanted SCT embryos [20]. Overexpressing *KDM4D* or *KDM4E* via mRNA injection into bovine SCT one-cell reconstructs also increased embryo development into high-quality blastocysts and survival into viable calves [21]. In pig, KDM4A mRNA injection improved *in vitro* but not in *in vivo* development [22]. In sheep, donor cell pretreatment with recombinant KDM4D protein improved *in vitro* development of SCT embryos [23] but *in vivo* development was not reported. A practical advantage of these mRNA- or protein-based approaches is that they are non-transgenic, which increases their applicability for agriculture.

Fourth, cellular quiescence by serum starvation (“G_0_“) has recently been shown to induce globally reduced DNA and histone trimethylation that correlate with a more relaxed G_0_ chromatin state and elevated somatic cell reprogrammability after SCT [24]. Specifically, H3K9me3 hypomethylation persisted in SCT-derived embryos and correlated with their increased survival into cloned cattle. Collectively, these findings established H3K9me3 as a major epigenetic barrier that obstructs restoration of totipotency following SCT.

Here we focused on the functional consequences of overexpressing murine *Kdm4b* in bovine embryonic fibroblasts (BEFs). We show a strong decrease in somatic H3K9me3 and H3K36me3 (H3K9/36me3) levels, which was rapidly restored in reconstructed bovine embryos after SCT. Sustained *Kdm4b* expression moderately increased reprogramming into cloned blastocysts. However, this treatment did not improve *in vivo* survival of cloned blastocysts into viable calves, indicating the need for alternative experimental strategies to robustly elevate cloning efficiency in livestock.

## MATERIALS AND METHODS

All animal studies were undertaken in compliance with the New Zealand Animal Welfare Act and were approved by the Ruakura Animal Ethics Committee.

### Vector construction

To generate the doxycycline (Dox)-inducible *Kdm4b-Egfp* piggybac (pB) transposon vectors, a pBS31 flp-in vector, kindly provided by J. Antony [15], was digested with *NotI* and *SalI* to excise the ∼2 kb *Kdm4b-Egfp* insert. This fragment comprises the 1-424 amino acid fully functional murine *Kdm4b*, fused to an N-terminal *Egfp* [25]. PB-TET-MKOS, a gift from Andras Nagy (Addgene plasmid # 20959), was modified to replace the original Tet-ON promoter [26] with a synthetic (GeneArt, Thermo Fisher, NZ) Dox-inducible P_TRE3G_ promoter that provides very low basal expression and high maximal expression after induction [27]. This pB-TRE3G response vector was further modified by replacing the β*geo* cassette with a *puromycin* selection marker, co-expressed with *Kdm4b-Egfp* via an internal ribosome entry site (IRES). Following *NotI*/*SalI* digest and cohesive end ligation, the newly generated *pB-TRE3G_Kdm4b-Egfp* vector was isolated using a PureLink™ HiPure Plasmid Filter Kit (Thermo Fisher, New Zealand) and validated by restriction enzyme digestion and sequencing analysis.

### Generation of stable BEF strains for inducible expression of *Kdm4b-Egfp*

As parental cells, a rejuvenated Dox-inducible driver line, based on BEFs carrying the Tet-On 3G transactivator driven by a human elongation factor 1 alpha (EF1α) promoter (Clonetech, Cat. No. 631167) was used (Green and Oback, manuscript in preparation) and induced with 2 µg/ml Dox (Sigma) two hours prior to transfection. At 70-80% confluence, these neomycin-resistant cells (‘BEF5-Tet’) were co-transfected with 2.5 μg *pB-TRE3G_Kdm4b-Egfp* using lipofection (Lipofectamine™ LTX/PLUS™, Thermo Fisher) and 2.5 μg pB transposase *CAGG-PBase* (pCyL43) [28]. Following selection with 1 µg/µl puromycin for seven days, 24 resistant cell clones were picked with cloning rings, subcloned onto gelatine-coated plates, expanded in puromycin-free medium, cryopreserved and two clones analyzed for Dox-inducible *Kdm4b-Egfp* expression.

### Induction of transgene expression

Both KDM4B-EGFP clones were cultured as described [29] in the presence of 2 µg/ml Dox for 24 hours. Induced cells were analyzed for EGFP fluorescence on a FACSCalibur™ (Becton and Dickinson, USA). To assess reversibility of KDM4B-EGFP expression, induced BEFs were washed, re-seeded in Dox-free medium and analyzed for EGFP fluorescence one day later. For analyses of serum-starved cells, fresh DOX was added to the low-serum cell culture medium every 2 days, while non-starved cells were induced for 2 days to prevent them from becoming confluent (Figs. 2-4, Tables 1-3).

**Table 1.** *In vitro* development of SCT embryos from induced vs non-induced *Kdm4b*-BEF donor cells.

| Donor treatment | Embryo treatment | n | N | No. of $\geq 1$-cells (% ± SEM) | No. of B1-3 (% ± SEM)<sup>+</sup> | No. B1-2 (% ± SEM)<sup>+</sup> |
| --- | --- | --- | --- | --- | --- | --- |
| NI | NI | 3 | 100 | 71 (71 ± 5%) | 59 (62 ± 2%) | 37 (37 ± 1%) |
| I | NI | 3 | 102 | 72 (71 ± 4%) | 58 (57 ± 3%) | 32 (31 ± 2%) |

Embryos were cloned from serum-starved donor cells (clonal strain #1) exposed to Dox (I) or buffer (NI) and cultured in medium without Dox (NI); n = number of independent SCT experiments; <sup>†</sup>percentage of embryos placed into IVC (N) that developed into D8 blastocysts (B) grade 1-3 (B<sup>1-3</sup>) or into B grade 1-2 (B<sup>1-2</sup>)

**Figure 1.**
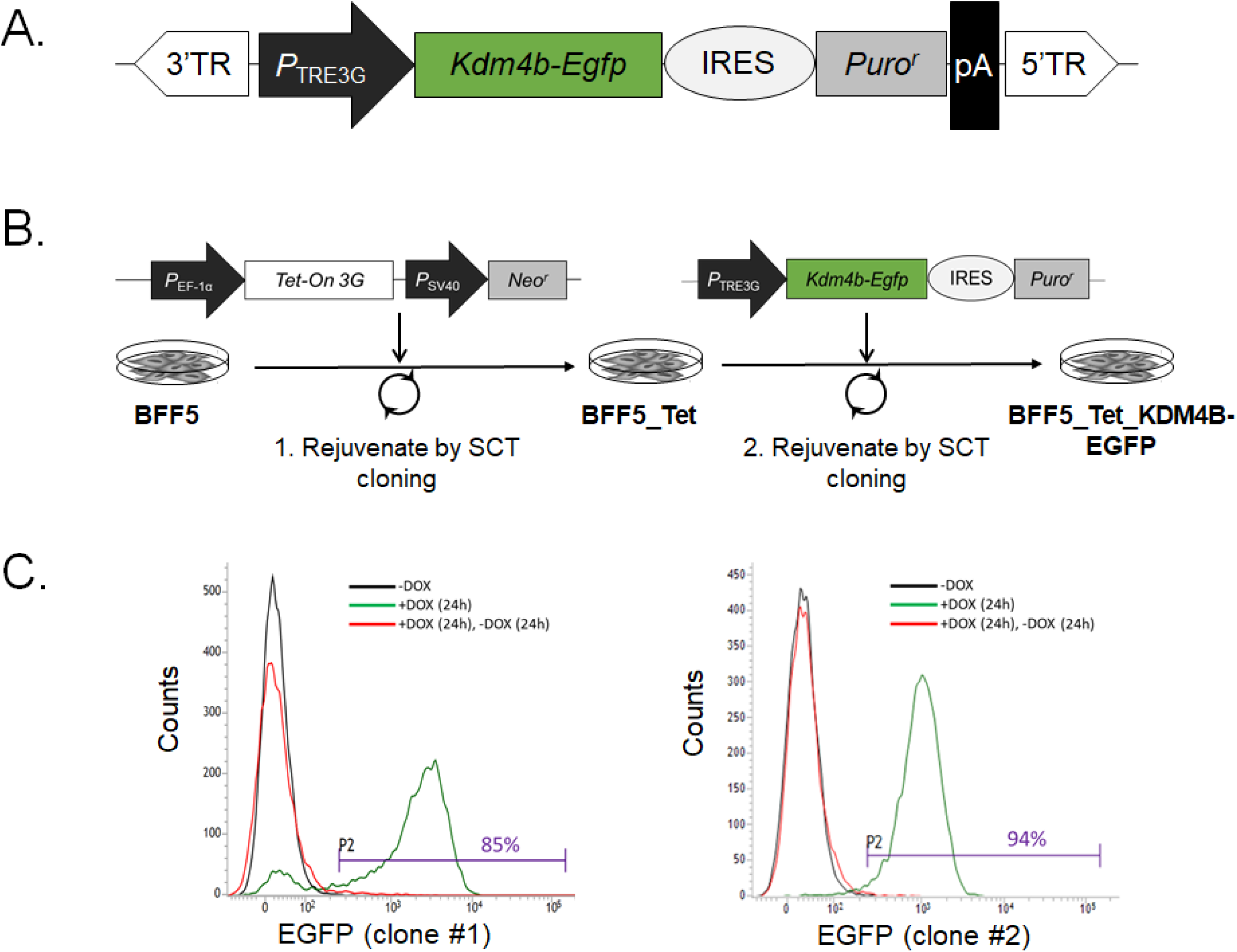
Dox-inducible Kdm4B-EGFP expression in rejuvenated bovine fibroblasts. **(A)** The piggyBac (PB) transposon vector used to deliver expression of tetracycline-responsive (TRE3G), puromycin-linked (IRES-puro-pA) histone demethylase (*Kdm4b-Egfp*). 3′/5′TR=pB terminal repeats; **(B)** Strategy for using somatic cell transfer (SCT) cloning to rejuvenate stable Dox-inducible BEFs that overexpress *Kdm4b-Egfp*. **(C)** Flow cytometry analyses of Kdm4b-EGFP induction in rejuvenated clonal BEF strains #1 (left) and #2 (right). Green fluorescence was determined using the FL1 EGFP emission channel. The range of intensities for green fluorescent cells (P2) is indicated. Relative cell number counts are plotted as a function of variable intensities of green fluorescence from individual cells. Line graph: non-induced (black line) vs induced (green line) cells; red line: induced cells one day after Dox-removal.

**Figure 2.**
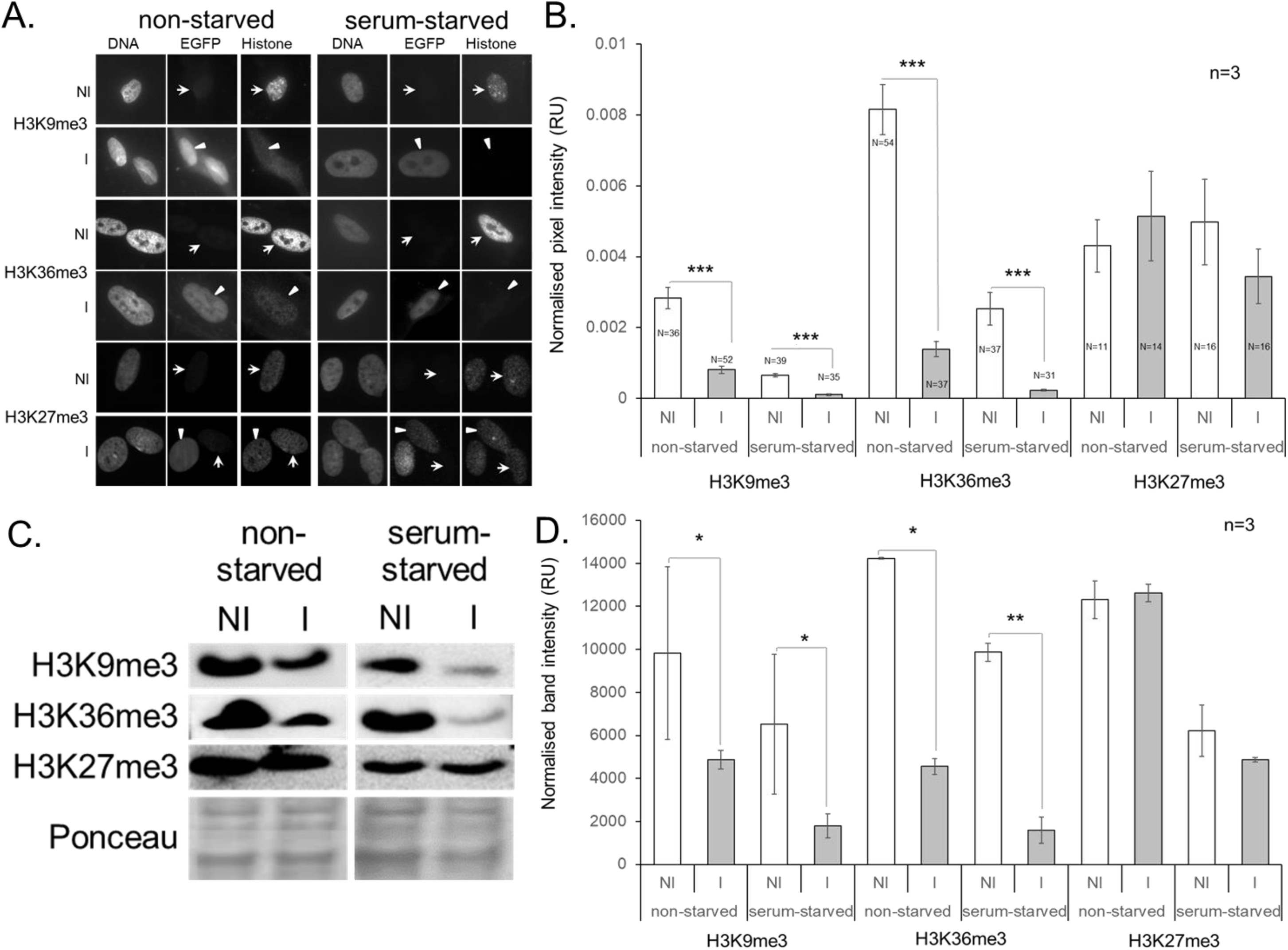
Kdm4B-dependant changes of H3Kme3 levels in BEFs. **(A)** Immunofluorescence analyses of different H3Kme3 modifications in non-induced (NI) and induced (I) *Kdm4B*-BEFs that were either serum-starved or non-starved. Cells were co-stained for DNA and with antibodies specific for EGFP and the indicated histone modification. Arrowheads and arrows indicate EGFP-positive and –negative nuclei, respectively, within the induced population. **(B**) Quantification of global H3Kme3 levels of indicated histone modifications from immunofluorescence images. Values represent normalized pixel intensity (region of interest/area) ± SEM. Asterisks indicate significant differences. N=number of nuclei quantified, n=3 biological replicates, RU=relative units; ***P<0.001; as determined by 2–tailed paired t-test. **(C)** Western analyses of histone extracts from induced (I) and non-induced (NI) *Kdm4b*-BEFs. Shown are representative immunoblots for three different methyl states of H3, normalized on three prominent invariant bands in the Ponceau stain. **(D)** Quantification of western analysis. Values represent normalized band intensity (histone modification/Ponceau) ± SEM determined from 3 biological repeats; RU=relative units; *, **: P<0.05 and P<0.01, respectively, as determined by 2–tailed paired t-test.

### RNA Extraction and PCR

Cells were lysed in 50 µL *RNA*GEM™ Tissue *PLUS* (containing 1.0 µL *RNA*GEM™) and cDNA synthesized as described [30]. Reverse transcriptase was omitted in one sample each time a batch was processed for cDNA synthesis (‘-RT’). Primers were designed using NCBI/ Primer-BLAST and synthesized by Integrated DNA Technologies (IDT, IA, USA). For quantitative PCR (qPCR), a LightCycler (Roche) was used with primers against transgene-derived *Kdm4b* (forward: 5’-AGAAGACACCGGGACCGATC-3’, reverse: 5’-TGAATTCATCCATGGTGGGG-3’, 190 bp amplicon), *ZFP37* (forward: 5’-CCTCAATGGGGTATCAGGCTC-3’, reverse: 5’-GATGGACTTCTTTTCCATTGGCT-3’, 156 bp amplicon) and *18S* (forward: 5’-AACGTCTGCCCTATCAACT-3’, reverse: 5’-AACCTCCGACTTTCGTTCT-3’, 70 bp amplicon).

All the reactions were performed with the KAPA SYBR® FAST qPCR Kit (Kapa Biosystems, MA, USA), which consisted of 0.4 µl of each primer (10 µM), 5.0 µl master mix, 2.2 µl DEPC water and 1.0-2.0 µl cDNA template. The following four-segment program was used: 1) denaturation (10 min at 95°C), 2) amplification and quantification (20 sec at 95°C, 20 sec at 55°C–60°C, followed by 20 sec at 72°C with a single fluorescent measurement repeated 45 times), 3) melting curve (95°C, then cooling to 65°C for 20 sec, followed by heating at 0.2°C/sec to 95°C while continuously measuring the fluorescence), and 4) cooling to 4°C. cDNA from bovine brain tissue provided a positive control. Product identity was confirmed by gel electrophoresis and melting curve analysis. For relative quantification, external standard curves were generated from serial 10-log dilutions for each gene in duplicate or triplicate. One high-efficiency curve (3.6 ≥ slope ≥ 3.1, R^2^ > 0.99) was saved and imported for relative quantification compared to 18S RNA as described [30]. A ‘no template’ control reaction was included in every run for each primer pair to test for DNA contamination and formation of primer–dimers.

### Quantitative Immunofluorescence (IF)

For each comparison, induced and non-induced cells were processed and analyzed in parallel. Cells were fixed and permeabilized simultaneously in 3.6% (w/v) paraformaldehyde/1% (w/v) Triton X-100 in PBS for 10 min, blocked with 2.5% (w/v) bovine serum albumin in PBS, all at room temperature, and incubated overnight at 4-8°C with the primary antibody specific for H3K9me3 (for immunofluorescence: rabbit polyclonal, gift by T. Jenuwein and previously validated for bovine [24]; for western analysis: mouse monoclonal, #c15200146, Diagenode), H3K36me3 (rabbit polyclonal, #9763, Cell Signaling) or H3K27me3 (mouse monoclonal, # c15200181, Diagenode). The next day, cells were washed in PBS-0.05% Tween 20 and incubated for 1 h at room temperature with Alexa Fluor® 488 or 568 donkey anti-goat, -mouse, -rabbit or –sheep secondary IgG antibodies (Thermo Fisher). DNA was counterstained with 5 µg/ml Hoechst 33342 (Sigma). Preparations were washed in PBS and H_2_O before mounting (DAKO, Med-Bio Ltd., New Zealand) onto glass slides. Wide-field epifluorescence (Olympus BX50) images were captured with a digital camera (Spot RT-KE slider) and Spot Basic software (v4.6).

SCT reconstructs were first fixed in 4% paraformaldehyde (30 min at 4°C) and then permeabilized in 0.1% Triton X-100 (5 min at room temperature) before blocking and immunostaining. Negative controls were processed the same way with blocking buffer instead of primary antibodies. Normalisation and quantification of global H3K9me3 levels was conducted in ImageJ. The Hoechst-stained region of interest (ROI) around the nucleus was outlined. Mean grey value intensity, measured at 3 random cytoplasmic locations, was subtracted from the mean ROI intensity. This background-corrected mean intensity represents the sum of grey values of all pixels in the ROI divided by the number of pixels and is referred to as normalized pixel intensity.

### Western blot

Histones were extracted using the EpiQuik™ Total histone extraction kit (Epigentek, Cat-OP-0006-100). Histone extracts (10-15 µg per lane) were separated on a 15% SDS PAGE gel, transferred onto a nitrocellulose membrane and probed with primary antibodies specified above, with the additional use of anti-H3K27me3 (rabbit polyclonal, #07-449, Millipore). Following incubation with a secondary antibody conjugated with horseradish peroxide, the modified histones were visualised with enhanced chemiluminescence. Signal intensity was normalized for three invariant histone bands (Ponceau S stain) and quantified using Quantity One software (Bio-Rad Laboratories Inc.).

### Generating SCT embryos and calves

Cultured *Kdm4b*-BEFs were serum-starved in medium containing 0.5% FCS for 3-6 days and harvested by trypsinization. Prior to each SCT run, cells were validated for successful induction of *Kdm4b* by monitoring EGFP signal with a digital fluorescence microscope (AMG-EVOS, Thermo Fisher). For IF analysis and embryo transfer (ET), zona-free SCT was performed as described [31]. Briefly, *in vitro* matured (IVM) non-activated metaphase II (MII)-arrested oocytes were derived from ovaries of slaughtered mature cows [31]. After IVM for 18-20 h, the cumulus-corona was dispersed by vortexing in bovine testicular hyaluronidase. Oocytes with a first polar body were chosen for enucleation. At 23-25 h post start of IVM, donor-cytoplast couplets were automatically aligned and electrically fused at 2.0 kV/cm. Reconstructed SCT embryos were artificially activated 3-4 h post-fusion, using a combination of ionomycin and 6-dimethylaminopurine (6-DMAP). After 4 h in 6-DMAP, reconstructs were washed three times in Hepes-buffered SOF (HSOF) and transferred into AgResearch-SOF single culture medium droplets for sequential *in vitro* culture (IVC). For inducing Kdm4b expression during IVC, embryos were placed into drops with fresh culture medium ± Dox on D0, D3 and D5. Embryo cultures were overlaid with mineral oil and kept in a humidified modular incubation chamber (ICN Biomedicals Inc., Aurora, OH) gassed with 5% CO_2_, 7% O_2_, and 88% N_2_. *In vitro* development was assessed on Day (D) 7 or D8 after fusion and morphological grade 1-2 quality embryos were non-surgically transferred singularly to synchronized recipient cows [31]. Fetal development was monitored by regular ultrasonography and rectal palpation throughout gestation.

### Statistical analysis

Values are the average of several replicates ± SEM. For the quantification of fluorescent signals and western blot bands, significance was determined by paired t-tests of normalized pixel intensities and expression ratios, respectively. For comparing *in vitro* development, significance was determined using the two-tailed Fisher exact test for independence in 2 X 2 tables. Significance was accepted as P<0.05. Unless stated otherwise, “N” denotes the number of samples analysed; “n” denotes the number of replicate experiments.

## RESULTS

### Rejuvenated bovine fibroblasts express Dox-inducible *Kdm4b*

To reduce H3K9/36me3 levels in bovine clones, we generated a piggyBac (PB) transposon vector for tetracycline-responsive (TRE3G) expression of a previously characterized murine *Kdm4b-EGFP* fusion construct [15], encompassing the fully functional catalytic KDM4B domain, translationally linked to a puromycin expression cassette (Fig. 1A). Using a two-step process of sequential rejuvenation (Fig. 1B), we first re-derived several BEFs carrying a constitutive *Tet-On 3G* transactivator construct (step 1), before transfecting these neomycin-resistant clonal cell strains (‘BEF5-Tet’) with *TRE3G_Kdm4b-Egfp* for a second round of rejuvenation (step 2). Detailed derivation and characterisation of selected BEF-Tet driver strains and inducible TRE-responder strains will be published elsewhere (Green and Oback, manuscript in preparation). To test induction of transgene expression, two rejuvenated *Kdm4b-Egfp* clonal BEF strains (#1, #2) were cultured in the presence of Dox for 24 hours. Due to the fusion with *Egfp*, cells expressing *Kdm4b* could be readily monitored and were analyzed by flow cytometry. An average 85% and 94% of *Kdm4b* strains #1 and #2, respectively, displayed EGFP fluorescence (Fig. 1C). We also examined the kinetics for switching off *Kdm4b-Egfp* expression. Within 24 hours after Dox removal, cells returned to a profile that was indistinguishable from non-induced cells, confirming that inducible transgene expression was fully reversible (Fig. 1C).

### KDM4B and serum starvation specifically reduce H3K9me3 and H3K36me3 in BEFs

We next evaluated the enzymatic activity of transgene-encoded murine KDM4B in both clonal BFF strains. Pairwise sequence alignment of bovine (XP_024850763.1) vs mouse (NP_742144.1) KDM4B showed high sequence conservation, with overall 87.7% identity (amino acids 1-424) and 100% identity within the JMJC domain (amino acids 176-292). To determine the effect on histone methylation, induced and non-induced *Kdm4b-Egfp* BEFs were analysed for levels of H3K9me3, and H3K36me3, as well as H3K27me3, which should not be directly targeted, by immunofluorescence and immunoblots. Induced cells expressing KDM4B-EGFP showed markedly reduced H3K9me3 and H3K36me levels (Fig. 2). In non-starved KDM4B-EGFP positive nuclei, signal intensity for H3K9me3 and H3K36me3 was 4-fold and 6-fold lower, respectively, than in non-induced controls (P<0.001), while non-targeted H3K27me3 control levels were not significantly changed (Fig. 2A, B). Following serum starvation and *Kdm4b* overexpression, signal intensity for H3K9me3 and H3K36me3 was 6-fold and 11-fold reduced, respectively, relative to non-induced controls (P<0.001), while H3K27me3 levels were again not significantly affected (Fig. 2A, B). Serum starvation has been shown to globally reduce trimethylation levels in bovine fibroblasts and increase their cloning efficiency [24]. We therefore analysed the effect of *Kdm4b* overexpression in non-starved vs serum-starved BEFs separately. Serum starvation alone reduced H3K9me3 levels 4-fold vs 8-fold (P<0.001) and H3K36me3 levels 3-fold vs 6-fold (P<0.001) in non-induced vs induced cells, respectively. Overall, Dox-induced, serum-starved *Kdm4b*-BEFs had the lowest H3K9me3 and H3K36me3 levels, 18-fold and 35-fold reduced, respectively, relative to non-induced, non-starved control BEFs.

To independently validate these results, we performed western blot analyses on bulk histones (Fig. 2C, D). For non-starved *Kdm4b-*BEFs, H3K9me3 and H3K36me3 levels dropped ∼2-fold (P=0.05) and ∼3-fold (P<0.05), respectively, compared to non-induced controls. Upon *Kdm4b* induction in serum-starved cells, H3K9me3 and H3K36me3 levels dropped ∼4-fold (P<0.05) and ∼6-fold (P<0.01), respectively, compared to non-induced controls. As for the IF characterization, Dox-induced, serum-starved *Kdm4b*-BEFs had the lowest H3K9me3 and H3K36me3 levels, while H3K27me3 levels were not significantly altered. For both induced and non-induced conditions, serum-starved BEFs showed lower normalized band intensities for all three histone trimethylations. Thus, both detection methods showed that BEFs were strongly and specifically de-trimethylated by serum starvation and inducible *Kdm4b-*overexpression.

### Up-regulation of *Zfp37* gene expression in *Kdm4*-BEFs

We previously surveyed the global transcriptional impact of *Kdm4b*-mediated reduction in H3K9/36me3 levels in female *Kdm4b-Egfp* MEFs by microarray, mRNA-seq and qPCR [16]. Across all three assays, the only consistently and significantly changed candidate transcript with a direct linkage to heterochromatin was *Zfp37*, which encodes a heterochromatin-associated zinc finger protein expressed in brain and testis [32]. Using RT-qPCR, Dox-induced, serum-starved donor BEFs showed 37-fold higher *Kdm4b* levels, resulting in a 4-fold up-regulation (P<0.05) of *Zfp37* (Fig. 3).

**Figure 3.**
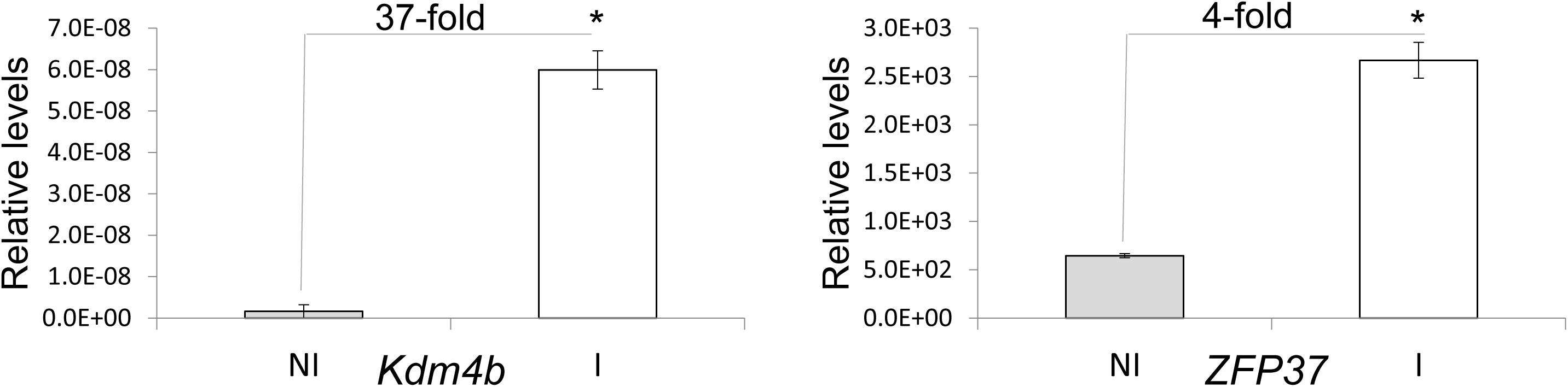
Kdm4B-dependant upregulation of *ZFP37* in serum-starved, induced BFFs. Shown are the relative expression levels of transgene-derived *Kdm4b* and *ZFP37*, relative to 18S expression (n=3 biological replicates). Results for non-induced (NI) and induced (I) *Kdm4b*-BEFs are represented by grey and white bars ± SEM, respectively, *=P<0.05.

### Reduced H3K9me3 and H3K36me3 donor cell levels are restored after SCT

We then examined how reduced H3K9/36me3 levels in serum-starved *Kdm4b*-induced BEFs changed after SCT and subsequent *in vitro* embryo development in the absence of Dox. SCT reconstructs from induced and non-induced *Kdm4b*-BEFs were fixed at various time points after electrofusion and analyzed by immunofluorescence for the presence of H3K9/36me3 (Fig. 4A). Ten minutes after fusing BEFs with the cytoplast, the EGFP signal had become undetectable in the SCT reconstruct, probably due to the high cytoplasmic dilution factor (data not shown). At this early time point, reduced H3K9/36me3 signals were still observed in SCT reconstructs from induced compared to non-induced donors (Fig. 4B). However, after three- and eight-hours post-fusion, the newly formed pseudo-pronuclei displayed similar H3K9/36me3 intensities in SCT reconstructs derived from induced and non-induced donor cells (Fig. 4B). No further differences in H3K9/36me3 staining intensities were observed at the two-cell (24 h) and D8 blastocyst stage (Fig. 4B). Thus, the initially reduced H3K9/36me3 levels in BEFs were restored relatively rapidly and induced hypomethylation did not persist after the late one-cell stage.

**Figure 4.**
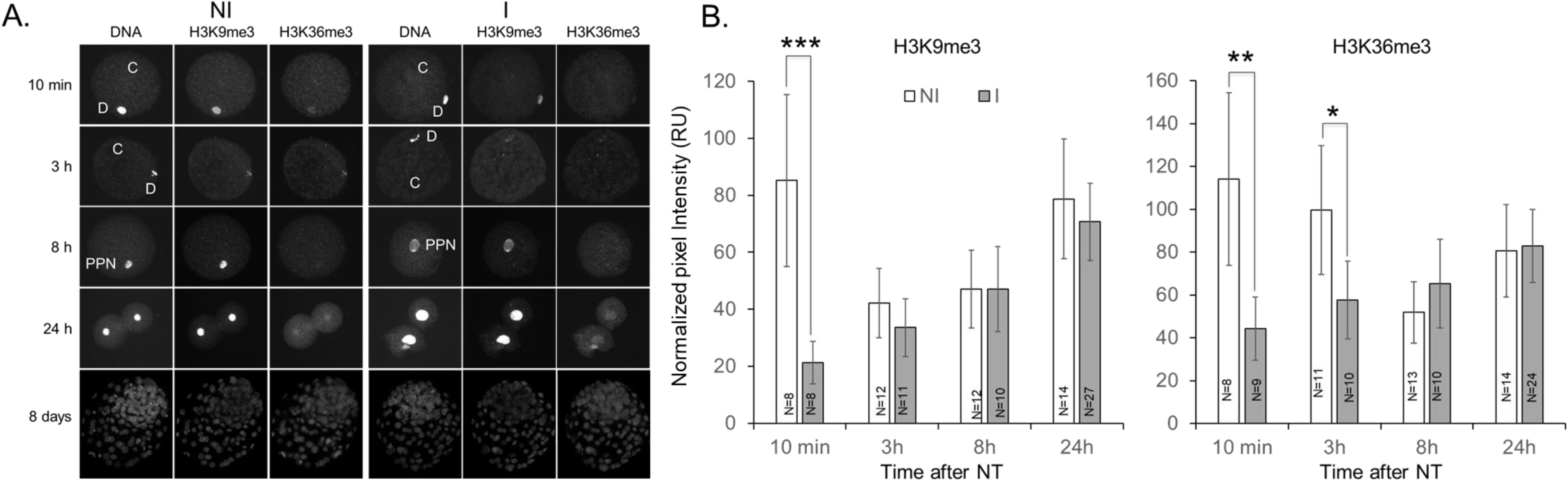
Characterization of global H3K9me3 and H3K36me3 levels following SCT with *KDM4b*-BEFs. SCT reconstructs were fixed and stained with H33342 (DNA) and antibodies specific for H3K9me3 and H3K36me3. **(A)** SCT reconstructs were analysed 10 minutes (upper panel), 3 h, 8 h, 24 h and 8 days (lower panel) after fusion of the donor cell with an enucleated oocyte (cytoplast). C= cytoplast; D=donor DNA; PPN=pseudo-pronucleus. **(B)** Quantification of immunofluorescence analysis in (A). Values represent normalized pixel intensity (region of interest/area) ± SEM. Asterisks indicate significant differences. N=number of NT reconstructs analysed. Asterisks indicate significant differences between BEFs from the same groups (induced or non-induced) at different time points, ***P<0.0001, ^**^P<0.01, ^*^P≤0.05 as determined by 2–tailed paired t-test.

### Sustained *Kdm4b* induction improves blastocyst development *in vitro*

To determine the effect of H3K9/36me3 hypomethylation on donor cell reprogrammability, we cultured SCT reconstructs to the blastocyst stage. Culture for embryos reconstructed with induced vs non-induced *Kdm4b-*BEFs was first performed without Dox, which showed no significant difference in total blastocyst development (31% vs. 37%, respectively, P=0.87, Table 1). We then continued induction during SCT embryo culture, refreshing Dox every two days, to determine the effect of continuous *Kdm4b* expression on development. Under these conditions, there was a significant improvement of total and a trend towards high-quality blastocyst development (58% vs. 42%, P<0.0001 and 29% vs 22%, P=0.057, respectively, Table 2) compared to non-induced controls.

**Table 2.**
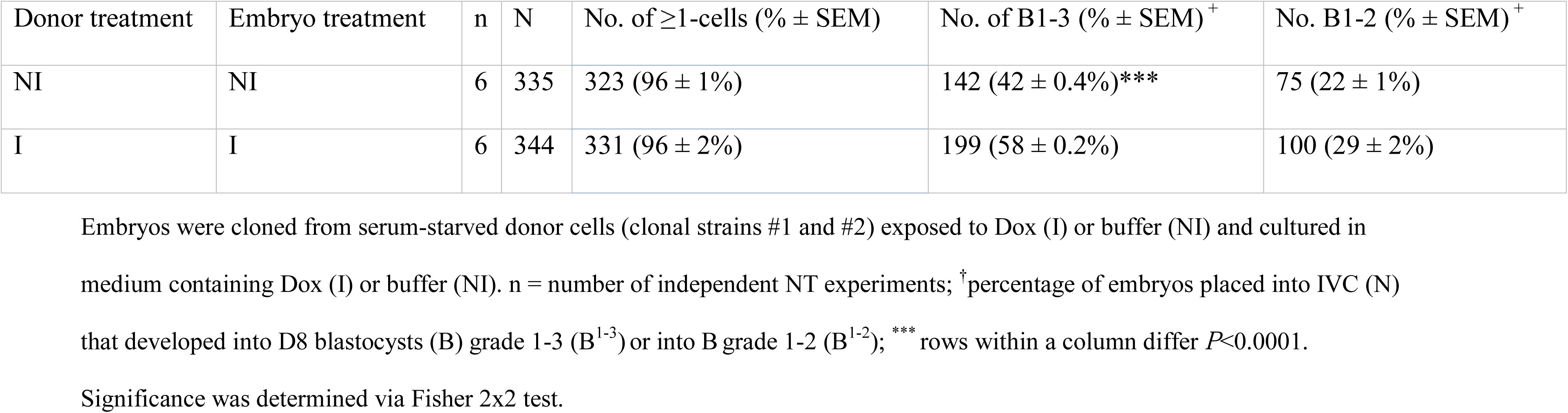
*In vitro* development of induced vs non-induced SCT embryos.

| Donor treatment | Embryo treatment | n | N | No. of $\geq 1$-cells (% ± SEM) | No. of B1-3 (% ± SEM)<sup>+</sup> | No. B1-2 (% ± SEM)<sup>+</sup> |
| --- | --- | --- | --- | --- | --- | --- |
| NI | NI | 6 | 335 | 323 (96 ± 1%) | 142 (42 ± 0.4%)*** | 75 (22 ± 1%) |
| I | I | 6 | 344 | 331 (96 ± 2%) | 199 (58 ± 0.2%) | 100 (29 ± 2%) |

Embryos were cloned from serum-starved donor cells (clonal strains #1 and #2) exposed to Dox (I) or buffer (NI) and cultured in medium containing Dox (I) or buffer (NI). n = number of independent NT experiments; <sup>†</sup>percentage of embryos placed into IVC (N) that developed into D8 blastocysts (B) grade 1-3 (B<sup>1-3</sup>) or into B grade 1-2 (B<sup>1-2</sup>); \*\*\* rows within a column differ $P < 0.0001$ . Significance was determined via Fisher 2x2 test.

### Reduced H3K9me3 and H3K36me3 donor cell levels do not improve fetal development *in vivo*

Long-term in utero development is a more stringent measure of donor cell reprogrammability into totipotency than blastocyst formation [33]. To assess the *in vivo* developmental potential of cloned embryos generated from *Kdm4b*-induced donor cells and embryos, which showed significantly improved blastocyst development compared to non-induced controls, D7 blastocysts were transferred into recipient cows. In three independent experiments, representing both *Kdm4b*-BEF strains, there were no significant differences for induced vs non-induced clones in establishing pregnancies and development to term (Table 3). One female calf, derived from non-induced *Kdm4b*-BEF donors, was delivered by Caesarean section on 18/01/2018. An ear biopsy was taken after two weeks and cultured ear skin fibroblasts Dox-treated for 48 hours, inducing nuclear KDM4B-EGFP signal in the re-derived cells (data not shown).

**Table 3.** *In vivo* development of induced vs non-induced SCT embryos.

| Donor group | Embryo group | n | N | D35 (% ± SEM)<sup>†</sup> | D60 (% ± SEM)<sup>†</sup> | D90 (% ± SEM)<sup>†</sup> | D120 (% ± SEM)<sup>†</sup> | D150 (% ± SEM)<sup>†</sup> | D180 (% ± SEM)<sup>†</sup> | D210 (% ± SEM)<sup>†</sup> | Term (% ± SEM)<sup>†</sup> | Adult (% ± SEM)<sup>†</sup> |
| --- | --- | --- | --- | --- | --- | --- | --- | --- | --- | --- | --- | --- |
| NI | NI | 3 | 32 | 12 (38 ± 6%) | 11 (34 ± 8%) | 4 (13 ± 8%) | 4 (13 ± 8%) | 3 (9 ± 8%) | 2 (6 ± 7%) | 2 (6 ± 4%) | 1 (3 ± 2%) | 1 (3 ± 2%) |
| I | I | 3 | 33 | 13 (39 ± 1%) | 8 (24 ± 10%)<sup>#</sup> | 3 (9 ± 7%) | 2 (6 ± 4%) | 2 (6 ± 4%) | 0 | 0 | 0 | 0 |

Embryos were cloned from serum-starved donor cells (clonal strains #1 and #2) exposed to Dox (I) or buffer (NI) and cultured in medium containing Dox (I) or buffer (NI). n = number of independent SCT experiments; <sup>†</sup>percentage of D8 blastocysts (N) that developed to this stage. <sup>#</sup>Two viable fetuses from the induced group were removed for cell line rejuvenation on D42 and D57, respectively, and numbers reduced accordingly. Potential survival of these fetuses would not have introduced significant differences between the two treatment groups.

Throughout gestation, we did not observe a difference in the frequency of hydroallantois, the most common complication with bovine SCT fetuses in our hands, which occurred twice in each treatment group (2/12 = 17% vs 2/13 = 15% for non-induced vs induced pregnancies, respectively). Thus, the moderate 1.4-fold increase in blastocyst reprogramming did not translate into an equivalent increase in post-blastocyst development.

## DISCUSSION

Here we show that the combination of *Kdm4b*-overexpression and serum starvation enhances simultaneous H3K9me3 and H3K36me3 removal from somatic donor cell chromatin. Following SCT, this hypomethylated epigenome reprograms better *in vitro* but not *in vivo*.

### Effects of H3K9me3/36me3 erasure on donor cell and embryo transcriptome

The beneficial effect of H3K9me3 removal on epigenetic reprogramming during SCT is now well described. A repressive modification, H3K9me3 causes heterochromatin formation [34] and epigenetic silencing [35]. Heterochromatinization generally marks megabase-scale gene-poor regions and repetitive elements, restricting access to chromatin-binding factors. Consequently, gene expression changes in Dox-induced somatic donor cells have been relatively minor [16]. Among the changed transcripts, the only direct linkage to heterochromatin was *Zfp37*. The encoded zinc finger protein is expressed in brain and testis, where it specifically associates with the heterochromatin adjoined to nucleoli [32]. Up-regulation of *Zfp37* was confirmed in *Kdm4b*-BEFs, supporting a conserved role of this target gene whose regulatory role is unknown. Overexpressing *Zfp37* in donor cells or embryos may shed light on its function and directly improve cloned blastocyst formation. However, similar attempts at rescuing development with *KDM4*-induced target genes, such as *Zscan4d* in mouse [18] and *SUPT4H1* in cattle [21], have failed. This highlights that a core gene regulatory network, rather than a single master regulatory transcription factor, may govern totipotency reprogramming.

In embryos, the donor cell transcriptome is largely repressed by the late one-cell stage in mouse, even though some highly expressed donor transcripts remain detectable for longer [18-22, 36, 37]. Exogenous expression of different KDM4 variants has normalized activation of developmentally regulated genes in SCT embryos, including *Kdm4b* (but not *Kdm4d*) and *Kdm5b* in mouse [17], *KDM4A* in pig [22] and *KDM1B, KDM4C, KDM4D* and *KDM4E* in cattle [21]. Some of these normalized genes lie within ‘reprogramming-resistant regions’ that fail to activate correctly at embryonic genome activation (EGA) in SCT embryos. These regions are conserved among different somatic cell types and species [18, 19]. They are relatively gene-poor and enriched for specific repeat sequences, such as LINE and LTR, but not SINE [18, 19]. Importantly, these regions are marked by H3K9me3 and enzymatic erasure of H3K9me3 facilitates their transcriptional activation, both for protein-coding genes and non-annotated repeat transcripts, restoring the global transcriptome of SCT embryos [18, 19, 21, 22]. Activation of repeat sequences around EGA is important for preimplantation development [38, 39]. In *Kdm4b*-expressing MEFs, endogenous retrovirus repeats, as well as major satellites, and intact LINEs, were derepressed [16]. Likewise, overexpression of *Kdm4d* and *KDM4E* partially relieved repression of endogenous retrotransposons and satellite I sequences in mouse [18] and bovine [21] embryos, respectively, around EGA.

In contrast to H3K9me3, it is presently unclear if and how reduced H3K36me3 levels affect reprogramming of the donor genome. H3K36me3 is present at the coding regions of transcribed genes [40]. Trimethylation of active H3K36 and repressive H3K27 is broadly mutually exclusive in euchromatin, which may prevent spreading and accumulation of silencing marks [40]. Indeed, global loss of H3K36me3 redistributes H3K27me2/3 from its endogenous sites to active gene bodies and mis-regulates gene expression [40]. Depleting H3K36me3 may have enriched for H3K27me3, for example, in regions maintaining X-linked gene expression and X chromosome inactivation in females [41]. These regions have been implicated in impairing SCT reprogramming through ectopic expression of *Xist* during the preimplantation period in both sexes, which can be corrected by removing or repressing *Xist* [13, 14, 22, 42].

### Serum starvation enhances H3K9/36me3 demethylation

We show that in serum-starved cells, histone trimethylation reduction for both H3K9me3 and H3K36me3 was about 2-fold greater under both non-induced and induced conditions. This 2-fold greater downregulation was observed using both IF and western blot analysis, even though the relative fold-changes were slightly different. It shows that serum starvation about doubles the effect of exogenously expressed *Kdm4b* on reducing histone trimethylation, confirming and extending previous results for H3K9me3 and H3K36me3 levels, respectively [24]. This additive effect of serum starvation suggests that it operates by a different molecular mechanism than induced *Kdm4b*.

We previously reported that serum starvation about halves the amount of H3K9me3 in non-transgenic primary bovine cell types, resulting in a more relaxed chromatin structure and providing a molecular correlate for the elevated reprogrammability of quiescent donor cells into totipotency. This halving may be due to starved cells acquiring non-methylated histones during S-phase but not re-instating trimethylation upon serum starvation, allowing completion of the last DNA replication and mitosis before entering a non-proliferative G_0_ state [24].

### Reduced H3K9/36me3 levels after SCT reprogramming

Globally reduced H3K9/36me3 levels in induced donor cells initially persisted in reconstructed embryos but returned to non-induced control levels within 3 hours after SCT. This kinetics is similar to the restoration of H3K9/36me3 marks previously observed for MEFs [16]. Metaphase-arrested mouse cytoplasts can also restore H3K9 methylation in transplanted male pronuclei within three hours [43]. The rapid gain of methylation at H3K9 sites suggests enzymatic action rather than slow incorporation of already trimethylated histones into newly synthesized DNA. At the 2-cell and blastocyst stage, we found no significant differences between SCT embryos derived from normal vs hypomethylated BEF donors, but we did not investigate any in-between stages, especially around EGA. Another immunofluorescence study of bovine SCT embryos has also found normal levels of H3K9me3 at the 2-4-cell and morula-blastocyst stage, however, signal intensity significantly increased at the 8-16 cell stage, indicating increased heterochromatin formation and gene silencing during EGA [21]. At this stage, bovine IVF embryos were also susceptible to *KDM4E* silencing, resulting in a further increase in H3K9me3 staining [21]. Pig SCT embryos that were injected with porcine *KDM4A* mRNA showed no change in global H3K9me3 levels at the 1-cell but significantly reduced levels at the 2-cell and blastocyst-stage [22]. This differs from our observations, which is likely due to the different time points of providing exogenous *Kdm4*. In our experiments, *Kdm4b* expression in the donor cells did not prevent re-methylation of the somatic genome within 3-24 hours after SCT. In the bovine and pig mRNA injection experiments, *Kdm4* expression started at the earliest 5 hr post-activation, corresponding to at least 5 h post-SCT, depending on the time interval between fusion and activation. Thus, H3K9me3 demethylation began at a time point when H3K9me3 re-methylation was already complete in our experiments. This delay in demethylation may be a conceptual advantage of mRNA injection over using hypomethylated donor cells. In all these studies, H3K9me3 levels were assessed by IF, which provides a read-out of global methylation levels. It cannot be excluded that certain somatic reprogramming-resistant genomic regions remained hypomethylated [18, 19]. Continued activation of locally H3K9/36me3-depleted regions may persist for some time, making the genes in these regions worthwhile candidates for further investigation in *Kdm4b*-induced vs non-induced SCT embryos at EGA. Thus, even a short-lived decrease in H3K9/36me3 levels could facilitate initial binding of oocyte reprogramming factors, triggering a ripple effect that persists after the global differences in H3K9me3/36me3 have disappeared. A similar effect was observed after knockdown of *Xist* in cloned male mouse morulae [13, 14]. Even though *Xist* expression returned to normal levels at the blastocyst stage, this transient *Xist* repression reactivated a number of X-linked genes in male cloned blastocysts and greatly improved their survival to term [13, 14].

### Kmd4-dependent hypomethylation and totipotency reprogramming

The discussed gene expression and epigenetic changes underlie the significantly improved blastocyst formation rate observed in all mRNA *Kdm4*-overexpression studies to-date, be it in donor cells [15, 16, 23] or in embryos [17-22]. However, blastocyst formation is a poor indicator for developmental competence [33]. Instead, development into healthy adult animals is the most definitive measure of extensive donor cell reprogramming [33]. *Kdm4b* expression in donor cells did not improve production of live calves. This is in contrast to another cattle study, which found pregnancy, birth and survival rates to be significantly higher in both *KDM4D*- and *KDM4E*-injected SCT embryos [18]. The overall increase in cloning efficiency by *KDM4D*- or *KDM4E*-injection was about 6-fold, an effect that we would have been able to detect statistically with the numbers of embryos transferred in our experiments.

The main differences between both studies are i) the variants used (*i.e.* C-terminally truncated murine *Kdm4b* vs full-length bovine *Kdm4D/E*) and ii) the method of *Kmd4* delivery (*i.e.* fusing multi-transgenic donor cells vs injecting mRNA into embryos). Truncated *Kdm4b* contained the JmjN domain, required for the activity of the JmjC catalytic center, but lacked the non-catalytic readers (Zn-finger PHD and double Tudor domains) that recognize specific histone lysine methylations [44]. These interactions target KDM4 to chromatin and regulate its activity, which is central to identify specific substrates and catalyse demethylation [44]. The KDM4B reader domain mediates exclusive binding to H3K23me3, stimulating the demethylase activity of full-length KDM4B towards H3K36me3 [45]. Deleting the C-terminal domain in KDM4 proteins can disrupt the cross-talk between reader and eraser domains, changing sub-cellular localization and demethylase activity, via altered chromatin affinity and/or protein complex formation with associated factors [46, 47]. However, we [15, 16] and others [25] showed that the PHD and Tudor domains are dispensable for reducing H3K9/36me3, but not H3K9me1/2 or H3K27me3, suggesting that truncated KDM4B would have achieved appropriate functionality in embryos. Similar to truncated KDM4B, both the PHD and Tudor domains are lacking in KDM4D/E. In contrast to full-length and truncated KDM4B, however, KDM4D/E has a different substrate specificity: it does not demethylate H3K36me3, yet attacks dimethylated in addition to trimethylated H1.4K26 [48, 49]. Furthermore, KDM4D demethylates H3K9me2 with similar efficiency as H3K9me3 and may even demethylate H3K9me1 [50, 51]. These differences may help to explain why KDM4D/E variants have improved cloning efficiency in mouse [18], macaque [20] and cattle [21], while studies with full-length KDM4A in pig [22] or truncated KDM4B in mice [15] have not. It is plausible that possessing H3K36me3 demethylase activity and/or lacking H3K9me1/2 and H1.4K26me3 demethylase activity interferes with long-term SCT-reprogramming success.

Different delivery approaches may have also contributed to the lack of a long-term beneficial effect in our experiments. In non-induced bovine embryos, *Kdm4b* expression and protein persistence would have likely been extinguished during the week it takes to reach the blastocyst stage. By contrast, mouse SCT reconstructs develop into blastocysts in about half the time. Hence the *Kdm4b-Egfp* transgene likely remained present for a relatively longer proportion of preimplantation development in mouse, especially during EGA. This could explain the significant boost in SCT blastocyst rates in mouse [16] but not cattle. It would be informative to measure the extent of donor-derived *Kdm4b* gene and protein expression in Dox-induced vs non-induced SCT embryos in more detail and compare its effect on H3K9/36me3 methylation patterns. Even in the presence of Dox, *Kdm4b-*expressing BEFs may not have maintained or activated full transcription of the transgene at the crucial time around EGA, which commences at the 8-cell stage in cattle [21], corresponding to about 3 days of *in vitro* culture. In that respect, mRNA injection post-activation may be more beneficial as it does not require transcription in a non-permissive environment prior to EGA but instead only relies on translation of the injected mRNA at the one-cell stage. In pig SCT, however, *KDM4A* mRNA injection into one-cells did not enhance *in vivo* cloning efficiency [22], suggesting that the *KDM4* variant plays a more important role than the delivery method. In this species, it was also noted that KDM4A mediated demethylation of the *XIST* promoter, resulting in *XIST* derepression [22]. Even though significantly elevated *XIST* expression was not observed with porcine KDM4B (or KDM4D), it would be important to investigate potential deregulation of *XIST* in female *Kdm4b*-BFFs and SCT embryos. If a similar phenomenon occurred with *Kdm4b* in cattle, this could have hindered long-term benefits in our experiments. To better distinguish variant vs delivery effects, *Kdm4d/e* bovine donor cells and truncated *Kdm4b* mRNA injection will need to be included in future comparisons.

In summary, we demonstrate that targeted reduction of repressive H3K9/36me3 marks (via *Kdm4b* overexpression), potentiated by global histone hypomethlation (via serum starvation), led to a derestricted genome with greater reprogrammability. Sustained *Kdm4b* overexpression in serum-starved BEFs and BEF-derived embryos improved *in vitro* reprogramming into cloned embryos. However, neither this cell-based approach, nor the injection method [22] have so far consistently improved epigenetic reprogramming into totipotency, emphasizing that KDM4-assisted SCT methods require empirical optimisation of the KDM4 variant and delivery method before being generally applicable to the cloning of livestock species. The surviving cloned cow conditionally overexpresses KDM4B, providing a new transgenic animal model to study the remodeling of heterochromatin architecture during differentiation and assess the role of H3K9/36me3 marks during nuclear reprogramming of different somatic and embryonic cell types.

## ACKNOWLEGEMENTS

We acknowledge Drs J. Antony, J. Li and A. Nagy for providing the *pBS31 flp*-in vector, *PB-TRE3G-Puro* plasmid backbone and *PB-TET-MKOS* plasmid (Addgene # 20959), respectively, as well as the Wellcome Trust Sanger Institute for making *PBase* (pCyL43) available. Dr. T. Jenuwein kindly donated anti-H3K9me3 antibody. We thank Jan Oliver for help with SCT cloning. This work was funded by the National Natural Science Foundation of China (NSFC, Grant No. 31460602; 31860645), the Ministry of Business, Innovation and Employment of New Zealand (contract C10X1002) and AgResearch.

